# CscoreTool: Fast Hi-C Compartment Analysis at High Resolution

**DOI:** 10.1101/188490

**Authors:** Xiaobin Zheng, Yixian Zheng

## Abstract

**Summary:** The chromosome conformation capture (Hi-C) has revealed that the eukaryotic genome can be partitioned into A and B compartments that have distinctive chromatin and transcription features. Current Principle Component Analyses (PCA)-based method for the prediction of A/B compartment prediction from Hi-C data requires substantial CPU time and memory. We report the development of a method, CscoreTool, that enables fast and memory-efficient determination of A/B compartments at high resolution even in dataset with low sequencing depth.

**Availability:** github.com/scoutzxb/CscoreTool

## 1 Introduction

The development of the proximity-ligation based methods for chromatin conformation capture (3C, 4C, 5C and Hi-C) has greatly improved the understanding of three-dimensional chromatin organization in the eukaryotic nucleus (Gibcus and Dekker, 2013). One important feature found in mammalian Hi-C studies is that the genome is organized into A or B compartments (Lieberman-Aiden, et al., 2009). Whereas the A compartment corresponds to genomic regions containing transcriptionally active and open chromatin (Lieberman-Aiden, et al., 2009), the B compartment corresponds to the peripheral heterochromatin associated with the nuclear lamina (van Steensel and Belmont, 2017). Recent studies showed that the A/B compartment organization is independent of the topologically associated domains (TADs) (Dixon, et al., 2012) and is more conserved than the organization of TADs at the single-cell level (Nora, et al., 2017; Stevens, et al., 2017). Therefore, understanding the A/B compartment organization is critical in deciphering 3D genome organization.

The current method for calculating A/B compartments is based on the Principal Component Analysis (PCA) of the normalized Hi-C interaction matrix (Lieberman-Aiden, et al., 2009). The first eigenvector (Principal Component 1, PC1) of the correlation matrix is defined as the compartment score, and genomic windows with positive or negative compartment scores are defined as A or B compartment, respectively. The PCA-based method has two major limitations. First, PCA is a descriptive statistical method designed for reducing dimension and the exact biological meaning of the compartment score is elusive. In this sense, compartment scores calculated from different Hi-C datasets may not be directly comparable. Second, PCA is slow and memory-inefficient when applied to large interaction matrix. This prohibits its application to high-resolution analysis of the compartment structure for most labs.

Here we proposed a statistical model to infer A/B compartments from Hi-C data. The output compartment score reflects the chance of a genomic window being in the A compartment. The implemented tool, namely CscoreTool, is ∼30 times faster and memory-efficient than existing PCA-based method for the same resolution. CscoreTool also works on high resolution for dataset with low sequencing depth.

## 2 Methods

We modeled the A/B compartments at individual cell level by allowing each individual cell to have its own A/B compartment segregation. Each genomic window *i* has a chance *P*_*i*_ to be in the A-compartment in an individual cell. Considering two genomic windows *i* and *j*, if their distributions in A and B compartments are independent within a cell population, the chance that they are in the same compartment status in an individual cell (defined as *F*_*ij*_) should be

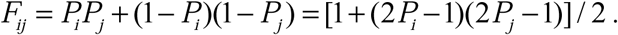

By defining C-score as *C*_*i*_ =2*P*_*i*_ −1, which ranges between −1 and 1, we rewrote *F*_*ij*_ =(1+*C*_*i*_ *C*_*j*_)/2.

As windows from different compartment status are unlikely to specifically interact with each other, the number of Hi-C contact between two genomic window *i* and *j* should be proportional to *F*_*ij*_. Considering the effect of linear genomic distance and experimental biases, we estimated the number of expected contact between windows *i* and *j* by *E*_*ij*_ = *B*_*i*_ *B*_*j*_ *H*(*d*_*ij*_) (1+*C*_*i*_ C_*j*_), where *d*_*ij*_ is the linear distance, *H*(*d*_*ij*_) is the function for distance dependency, *B*_*i*_ and *B*_*j*_ are the bias factors from Hi-C experiments. We assumed that the observed number of contact (*n*_*ij*_) follows a Poisson distribution *n*_*ij*_ ∼ Poisson(*E*_*ij*_), and deduced a log-likelihood function

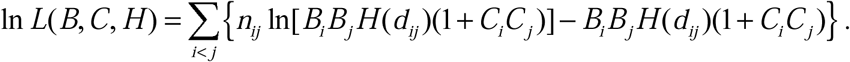

Where *d*_*ij*_ is the linear distance, *H*(*d*_*ij*_) is the function of the distance dependency, *B*_*i*_ and *B*_*j*_ are the bias factors from Hi-C experiment, and *C*_*i*_ and *C*_*j*_ are the C-scores.

Here since we assumed the distribution of window *i* and *j* in A and B compartments are independent with each other, we used only window pairs with at least 1Mb distance. This is an adjustable parameter in the CscoreTool with 1Mb as the default. We also used only intra-chromosome interactions to reduce noise. We then used a maximum-likelihood approach to estimate the C-scores.

Note that - ln *L* is a convex function for each individual parameter. Thus our maximum-likelihood estimation was done by iteratively optimizing each of the model parameter one by one conditioning on other parameters, which ensures monotone increase of the likelihood function in each step. This monotone increase of the likelihood function ensures convergence based on the monotone convergence theorem. Details of the optimization steps are described as below:

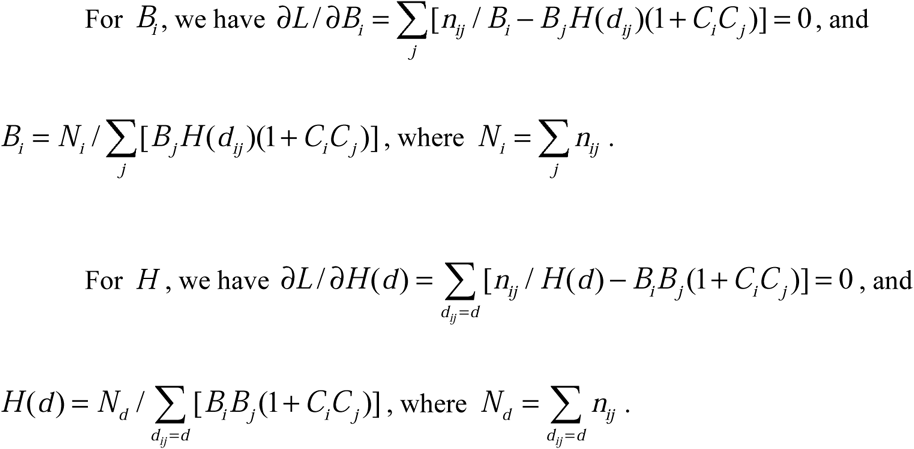

Since *N*_*d*_ may be low for large *d*, we used exponentially growing distance bins [*d*_*n*_, *d*_*n* +1_), where *d*_0_ = 1 Kb, each *d*_*n*+1_ = *d*_n_ × 10^0.04^, and all distances within each bin were regarded as equal.

For *C*_*i*_, we remove irrelevant parts in the log-likelihood function and get ln 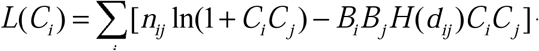 + *constant.* The maximum point of this function in (-1,1) does not have an explicit solution, but can be solved by Newton method, since - ln *L* is a convex function of *C*_*i*_.

The whole algorithm works as following: We first randomly generated the initial parameters, and then we performed iterative optimization for each parameter one by one as described above, in the order of *H*, *B* and *C.* The algorithm is considered convergent when the increase of the value of the likelihood function between two rounds (one round means updating all parameters once) is less than 1, and we can obtain the convergent values of C-scores. Each chromosome was analyzed separately, since inter chromosome reads were not used. In practice, the algorithm converged in ∼ 20 rounds, and was not sensitive to the initial parameters, since we tried 10 random initial parameters and obtained the same results. We implemented the algorithm with C++ and tested the resultant tool, namely CscoreTool, on the high resolution Hi-C dataset of the GM12878 cell line (Rao, et al., 2014). Among PCA-based variants, we chose HOMER for comparison (Heinz, et al., 2010), as it uses the computing-efficient R language (implemented by C and Fortran) for PCA. Mapped reads (to mm9) for Hi-C libraries HIC001-HIC029 were downloaded from the GEO database (GSE63525). Only reads with MAPQ ≥ 30 at both ends were kept, and since the analyses for each chromosome are independent, we focused our test on chromosome 1. CscoreTool was tested for 1Kb, 5Kb, 10Kb, 25Kb, 50Kb and 100Kb resolutions, while HOMER (runHiCpca.pl) was tested for 10Kb, 25Kb, 50Kb and 100Kb resolutions with all the other parameters set as default. When testing HOMER at a resolution of 5Kb, we obtained a warning message from HOMER indicating that the matrix was too big to handle. All tests were performed on the Memex high-performance computer of Carnegie Institution for Science. Although CscoreTool supports parallelization, we used only one CPU (2.5GHz) for the comparison, since HOMER does not support parallelization within one chromosome.

## 3 Results

We first compared the running time and memory usage between CscoreTool and HOMER. At the same resolutions, CscoreTool is over 30 times faster than HOMER (Fig. 1A). HOMER uses 40 min-3 h for PCA analysis at 25-100Kb resolutions but requires over 20 hours for 10Kb-resolution, and it stopped running for 5Kb resolution because it could not handle the large matrix. By contrast, CscoreTool used a few minutes for analyses at 25-100Kb resolution; 35 min for 10Kb resolution; 2 hours for analyses at 5Kb resolution; and ∼3 days for analysis at 1Kb resolution. The time consumed can be further reduced by using parallelization. For example, using 12 CPUs, the 1Kb-resolution analysis can be done in ∼12 hours. The memory usage of CscoreTool is also much less than HOMER (Fig. 1B). The 1Kb-resolution analysis by CscoreTool used less than 10GB memory, whereas HOMER used more than 60GB memory for the same dataset at the 10-100Kb resolutions that we could test. Thus, CscoreTool is much faster and more memory-efficient than the PCA-based compartment analysis method.

**Figure 1.**
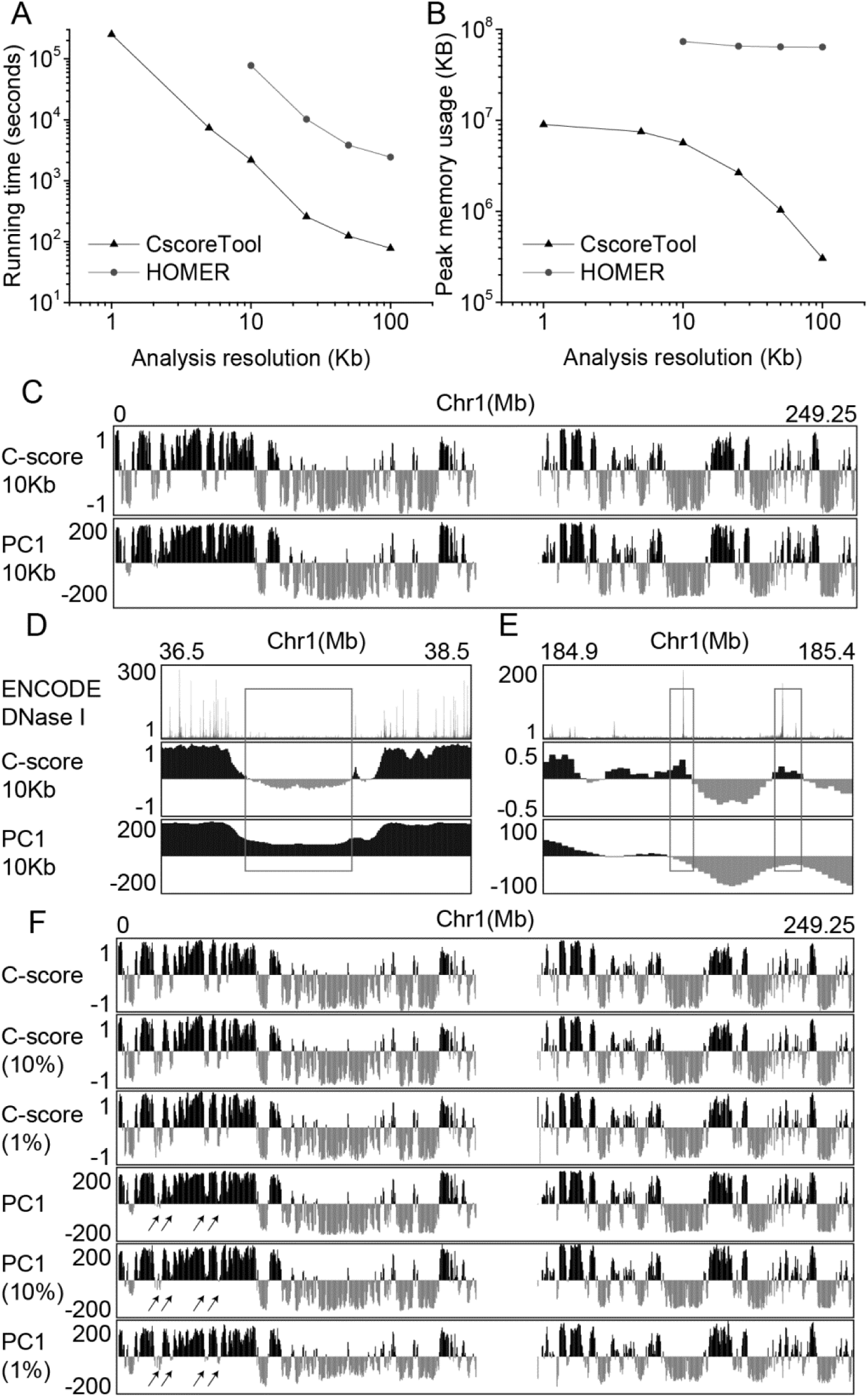
**A-B**. Comparison of running time and memory usage between CscoreTool and HOMER. **C**. Whole-chromosome view of C-score by CscoreTool and PC1 by HOMER calculated at 10Kb resolution. **D-E**. Example regions showing CscoreTool can detect smaller B-or A-compartments missed by HOMER (in rectangles). **F**. Comparison of the results by CscoreTool and HOMER at 10Kb resolution when only small fractions (10% or 1%) of the reads are used. Arrows indicate regions showing inconsistencies among different resolutions analyzed by HOMER.

We then compared the distribution of C-score from CscoreTool to the PC1 score from HOMER at 10Kb resolution. Whole-chromosome view showed that the patterns are in general similar (Fig. 1C), indicating that both scores capture the large-scale structure of A/B compartments. Finer scale comparison reveals some regions showing differences between the methods (Fig. 1D-E). For example, HOMER predicted a long chromatin region as compartment A (Fig. 1D), whereas CscoreTool showed that a long stretch of chromatin within this region belonged to compartment B. Since DNaseI sensitivity is associated with chromatin in compartment A (Lieberman-Aiden, et al., 2009), we analyzed the ENCODE DNase I data (Consortium, 2012) in these cells and found that the compartment B chromatin predicted by CscoreTool had a very low DNaseI sensitivity (Fig. 1D), indicating that our method correctly predicted this region as B compartment. We also found small A-compartments with DNase I peaks that were missed by HOMER (Fig. 1E). These results show that CscoreTool is better at detecting smaller A-and B-compartments than HOMER.

The high-resolution data we used here have 3.51G mapped non-redundant reads, which requires substantial amount of sequencing that are not practical for many labs and not necessary for many Hi-C experiments. Therefore, we created two smaller datasets by randomly selecting 10% and 1% of from the 3.51G mapped reads. The 10% dataset (mid-depth) corresponds to 351M mapped non-redundant reads, which is common for most Hi-C read depth. The 1% dataset (low-depth) corresponds to 35.1M mapped non-redundant reads, which is common for low-cell-number Hi-C data such as those in early embryo development (Du, et al., 2017; Ke, et al., 2017). We then tested the performance of CscoreTool and HOMER on these two smaller datasets. We found that CscoreTool gave very consistent results among all sequencing depth (Fig. 1F) on large scale. By contrast, HOMER showed more inconsistency between different analysis resolutions (Fig. 1F). Taken together, we show that CscoreTool can perform accurate high-resolution compartment analysis at both high and low sequencing depth with similar accuracy and it requires less time and computer memory than the commonly used PCA-based methods.

## References

Consortium, E.P. (2012) An integrated encyclopedia of DNA elements in the human genome, Nature, 489, 57–74.

Dixon, J.R., et al. (2012) Topological domains in mammalian genomes identified by analysis of chromatin interactions, Nature, 485, 376–380.

Du, Z., et al. (2017) Allelic reprogramming of 3D chromatin architecture during early mammalian development, Nature, 547, 232–235.

Gibcus, J.H. and Dekker, J. (2013) The hierarchy of the 3D genome, Molecular Cell, 49, 773–782.

Heinz, S., et al. (2010) Simple combinations of lineage-determining transcription factors prime cis-regulatory elements required for macrophage and B cell identities, Molecular Cell, 38, 576–589.

Ke, Y., et al. (2017) 3D Chromatin Structures of Mature Gametes and Structural Reprogramming during Mammalian Embryogenesis, Cell, 170, 367–381 e320.

Lieberman-Aiden, E., et al. (2009) Comprehensive mapping of long-range interactions reveals folding principles of the human genome, Science, 326, 289–293.

Nora, E.P., et al. (2017) Targeted Degradation of CTCF Decouples Local Insulation of Chromosome Domains from Genomic Compartmentalization, Cell, 169, 930–944 e922.

Rao, S.S., et al. (2014) A 3D map of the human genome at kilobase resolution reveals principles of chromatin looping, Cell, 159, 1665–1680.

Stevens, T.J., et al. (2017) 3D structures of individual mammalian genomes studied by single-cell Hi-C, Nature, 544, 59–64.

van Steensel, B. and Belmont, A.S. (2017) Lamina-Associated Domains: Links with Chromosome Architecture, Heterochromatin, and Gene Repression, Cell, 169, 780–791.

